# A rapid and robust leaf ablation method to visualize bundle sheath cell chloroplasts in C_3_ species

**DOI:** 10.1101/2023.03.31.535114

**Authors:** Kumari Billakurthi, Julian M Hibberd

## Abstract

**Background:** It has been proposed that engineering the C_4_ photosynthetic pathway into C_3_ crops could significantly increase yield. This goal requires an increase in the chloroplast compartment of bundle sheath cells in C_3_ species. To facilitate large-scale testing of candidate regulators of chloroplast development in the rice bundle sheath, a simple and robust method to phenotype this tissue in C_3_ species is required.

**Results:** We established a leaf ablation method to accelerate phenotyping of rice bundle sheath cells. The approach allowed bundle sheath cell dimensions, chloroplast area and chloroplast number per cell to be measured. Using this method, bundle sheath cell dimensions of maize were also measured and compared with rice. Our data show that bundle sheath width but not length significantly differed between C_3_ rice and C_4_ maize. Comparison of paradermal versus transverse bundle sheath cell width indicated that bundle sheath cells were intact after leaf ablation. Moreover, comparisons of planar chloroplast areas and chloroplast numbers per bundle sheath cell between wild-type and transgenic rice lines expressing the maize *GOLDEN-2* (*ZmG2*) showed that the leaf ablation method allowed differences in chloroplast parameters to be detected.

**Conclusions:** Leaf ablation is a simple approach to accessing bundle sheath cell files in C_3_ species. We show that this method is suitable for obtaining parameters associated with bundle sheath cell size, chloroplast area and chloroplast number per cell.

## Background

Photosynthesis is fundamental to life on earth and allows assimilation of atmospheric CO_2_ into biomass via the Calvin-Benson-Bassham or C_3_ cycle [1–3]. In plants the photosynthetic process is broadly categorised into C_3_, C_4_ and Crassulacean Acid Metabolism based on the pathway of carbon fixation. However, plants that use C_3_ photosynthesis predominate such that species using C_4_ and Crassulacean Acid Metabolism account for only three and six percent of land plants respectively [4–6]. In C_3_ plants, mesophyll cells are filled with chloroplasts and so are the major site of photosynthesis (Figure 1a). In these plants the enzyme Ribulose-1,5-Bisphosphate Carboxylase/Oxygenase (RuBisCO) carboxylates the five-carbon compound Ribulose-1,5-bisphosphate (RubP) via the C_3_ cycle to generate two molecules of the three-carbon compound 3-phosphoglycerate. In contrast, in the vast majority of C_4_ plants the reactions of carbon assimilation are equally partitioned between mesophyll and bundle sheath cells. HCO_3_^-1^ is initially fixed in mesophyll cells by Phospho*enol*pyruvate Carboxylase (PEPC) to generate four-carbon compounds such as malate and aspartate that then diffuse into bundle sheath cells. Decarboxylation of either aspartate or malate in the bundle sheath releases high concentrations of CO_2_ in bundle sheath cells that can then be assimilated by RuBisCO [7].

**Figure 1:**
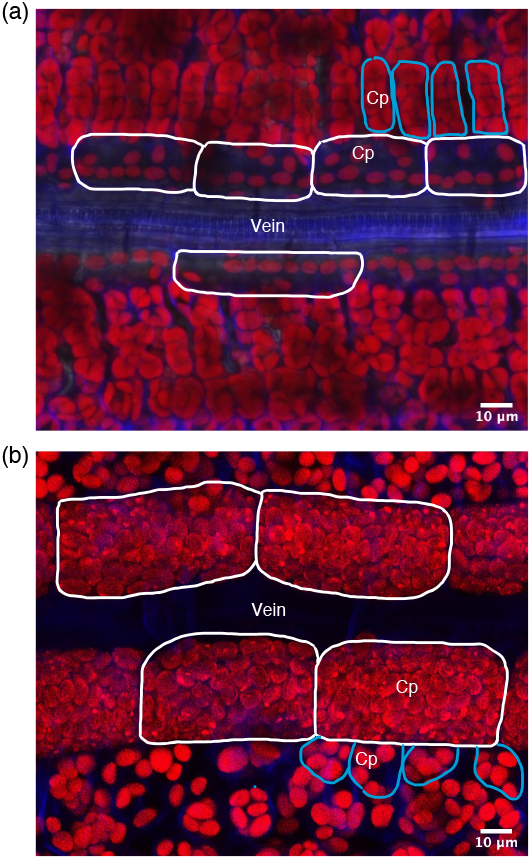
Confocal laser scanning microscopy images of C_3_ (rice) and C_4_ (maize) mesophyll and bundle sheath cells. Images are derived from paradermal sections. Representative maximum intensity projection image of a Z-stack from wild type rice (a) and maize (b). Bundle sheath and mesophyll cells are highlighted with white and blue lines respectively. Chloroplasts from bundle sheath cells of maize generate lower autofluorescence due to lower amounts of Photosystem II. Cp: Chloroplasts.

Due to the C_4_ cycle concentrating CO_2_ around RuBisCO, C_4_ species are more efficient under dry and high-temperature conditions. Moreover, they often have improved water and nitrogen use efficiencies compared with C_3_ plants [8–11]. Apart from those species that use single-celled C_4_ photosynthesis [12], a unifying character underpinning the C_4_ pathway is a specialised form of leaf morphology termed Kranz anatomy [13]. Kranz anatomy is characterised by a high vein density and bundle sheath cells that are altered both morphologically but also in terms of organelle occupancy and positioning. During the C_3_ to C_4_ trajectory, evolution has generated bundle sheath cells that are larger in the medio-lateral leaf axis [14,15] and contain numerous larger chloroplasts ([16], Figure 1).

Increasing the photosynthetic efficiency of C_3_ crops would help meet future demands for food, especially under changing climatic conditions. It has been predicted that introducing the C_4_ pathway into C_3_ crops could increase their photosynthetic efficiency by up to 50 % [17]. However, one of the main bottlenecks is an incomplete understanding of how bundle sheath cells become photosynthetically activated in C_4_ plants. On average, the bundle sheath chloroplast content of C_4_ species is ∼ 30 % more than in C_3_ species [16,18], but how this evolved is not fully understood. The GOLDEN2-LIKE family of transcription factors known to regulate chloroplast development in C_4_ species [19–21]. Although overexpression of *GOLDEN2* or *GOLDEN2-LIKE 1* from C_4_ *Zea mays* in rice increased bundle sheath chloroplast volume, this did not phenocopy the increase in chloroplast occupancy found in C_4_ plants [22].

Introducing C_4_ bundle sheath anatomy into C_3_ rice is therefore likely to involve large-scale testing of candidate genes involved in bundle sheath cell and chloroplast development and phenotyping bundle sheath cells. However, the bundle sheath has been challenging to phenotype in C_3_ plants. Classical bright-field light microscopy after embedding samples in resin and thin sectioning has been used [18]. Although, this is simple and easily available, it only captures two-dimensional (2D) information from a thin section. 2D-transmission electron microscopy (2D-TEM) is widely used for characterising the ultracellular structure and organisation in photosynthetic cell types [23] but it is expensive and has the same limitations as light microscopy when cell and chloroplast parameters are being quantified. A single-cell isolation method has been established to study mesophyll and bundle-sheath cell dimensions and chloroplast occupancy, but it requires enzymatic digestion of leaf tissue that might disturb cell integrity and chloroplast size [22]. Lastly, more advanced electron microscopy-based 3D reconstruction methods such as serial block-face scanning electron microscopy (SBF-SEM) can cover large fields of view and reconstruct ultrastructural features in 3D such that volume of leaf cells and chloroplasts can be quantified [24]. However, it is costly and labour-intensive. Thus, each of these approaches has disadvantages for high-throughput screening of bundle sheath cells in species such as C_3_ rice.

To address this, we established a simple and robust method to expose bundle sheath cell files in rice and measure their cell dimensions, as well as the chloroplast area and chloroplast number per cell. We show that these bundle sheath cells are intact and the chloroplast number per cell is comparable with previous reports [22]. We also applied this method to the C_4_ species maize to measure bundle sheath cell dimensions and made comparisons between bundle sheath cells in these two species. When combined with genetic perturbations we anticipate that this approach will provide insight into structure function relations of bundle sheath cells in species such as rice.

## Results

### A simple and robust method to visualize bundle sheath cells in C_3_ rice and C_4_ maize

The middle region of fully expanded fourth leaves from rice and maize was fixed with glutaraldehyde. Prior to ablation, although parallel venation was detectable in rice at low magnification, when higher power objectives were used the significant amount of light scattering meant that individual cells including the bundle sheath were not visible (Figure 2a&c). However, bundle sheath strands and cells became visible (Figure 2b&d) after the adaxial side of leaves was ablated by gentle scraping (Additional file 1). In rice scraping was carried out until mesophyll tissue surrounding intermediate veins appeared less green. As the bundle sheath is deep in the C_3_ leaf because of the many layers of mesophyll cells [25], two to three minutes of ablation (Additional file 1) was required to expose bundle sheath cells around intermediate veins (Figure 2b&d). Consistent with rice leaf anatomy, three to four intermediate veins (rank-1; tertiary; 3°) were present between the larger lateral (secondary; 2°) veins.

**Figure 2:**
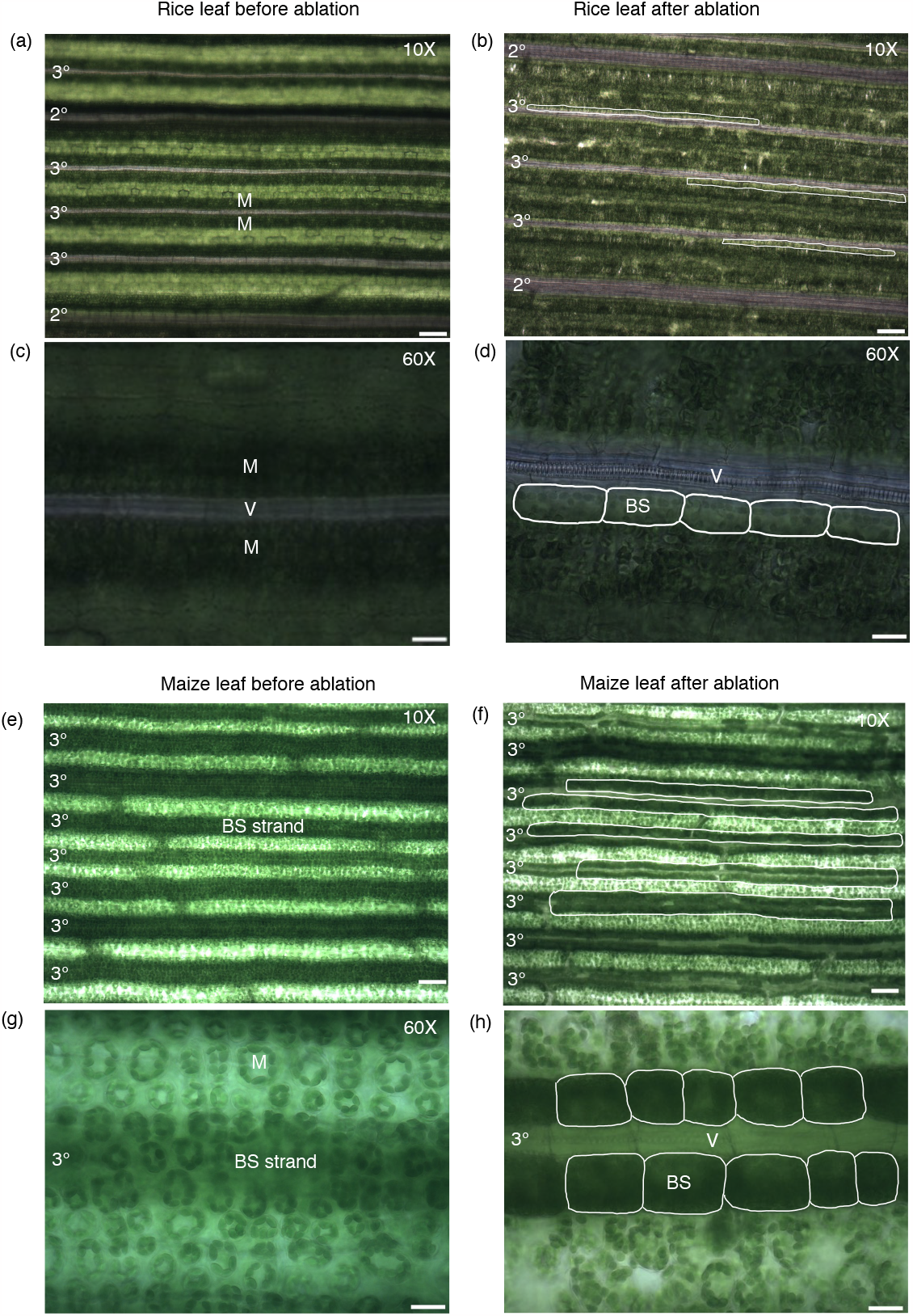
Visualization of rice and maize leaves before and after ablation. Light microscopy images of rice (a-d) and maize (e-h) leaves before (left) and after scraping (right). Low power images (b, f) illustrating the impact of ablation on bundle sheath visibility (highlighted with white lines). Representative images of selected regions at higher magnification (d, h). Bundle sheath cells highlighted with white lines. Abbreviations are as follows: 2°: secondary veins; 3°: tertiary veins; V: Vein; BS: Bundle sheath cell. M: Mesophyll tissue. Scale bar represents 100 μm (a, b, e, f) and 20 μm (c, d, g, h).

In maize, dark green strands that represent the bundle sheath were visible prior to scraping (Figure 2e) and although mesophyll cells were detectable at higher magnification this was not true for the bundle sheath (Figure 2g). Scraping of maize allowed files of dark green bundle sheath and the less green mesophyll cells to be identified (Figure 2f). C_4_ maize has increased numbers of intermediate (rank-1 + rank-2) veins between the larger laterals because of an increase in the density of rank-2 intermediates [26] and leaf ablation was consistent with this (Figure 2f). In C_4_ maize it took less than one minute to ablate mesophyll layers such that bundle sheath cell files were clearly visible (Figure 2f&h).

### Quantification of bundle sheath cell dimensions

To provide quantitative insight into differences between bundle sheath cells of C_3_ rice and C_4_ maize we ablated leaf tissue from each species and then used calcofluor white to mark cell walls (Figure 3a&b). Bundle sheath cell length and width measurements were taken at the mid-point of both the proximal-distal and medio-lateral axes, and cell area was calculated (Figure 3a&b). Average bundle sheath cell width was 15 μm in rice and 32 μm in maize (Figure 3c) but there is a very small difference in bundle sheath cell length between these two species (Figure 3d). However, as a consequence of the increased width of bundle sheath cells in maize, mean bundle sheath cell area was significantly higher (1404 μm^2^) than that of rice (598 μm^2^) (Figure 3e). We also observed high variance in bundle sheath cell dimensions in both rice and maize. This variation in width and length of bundle sheath cells was two-fold and three-fold respectively in both species (Figure 3c&d), and the variation in bundle sheath cell area of maize was around 1.4 times greater than that of rice (Figure 3e).

**Figure 3:**
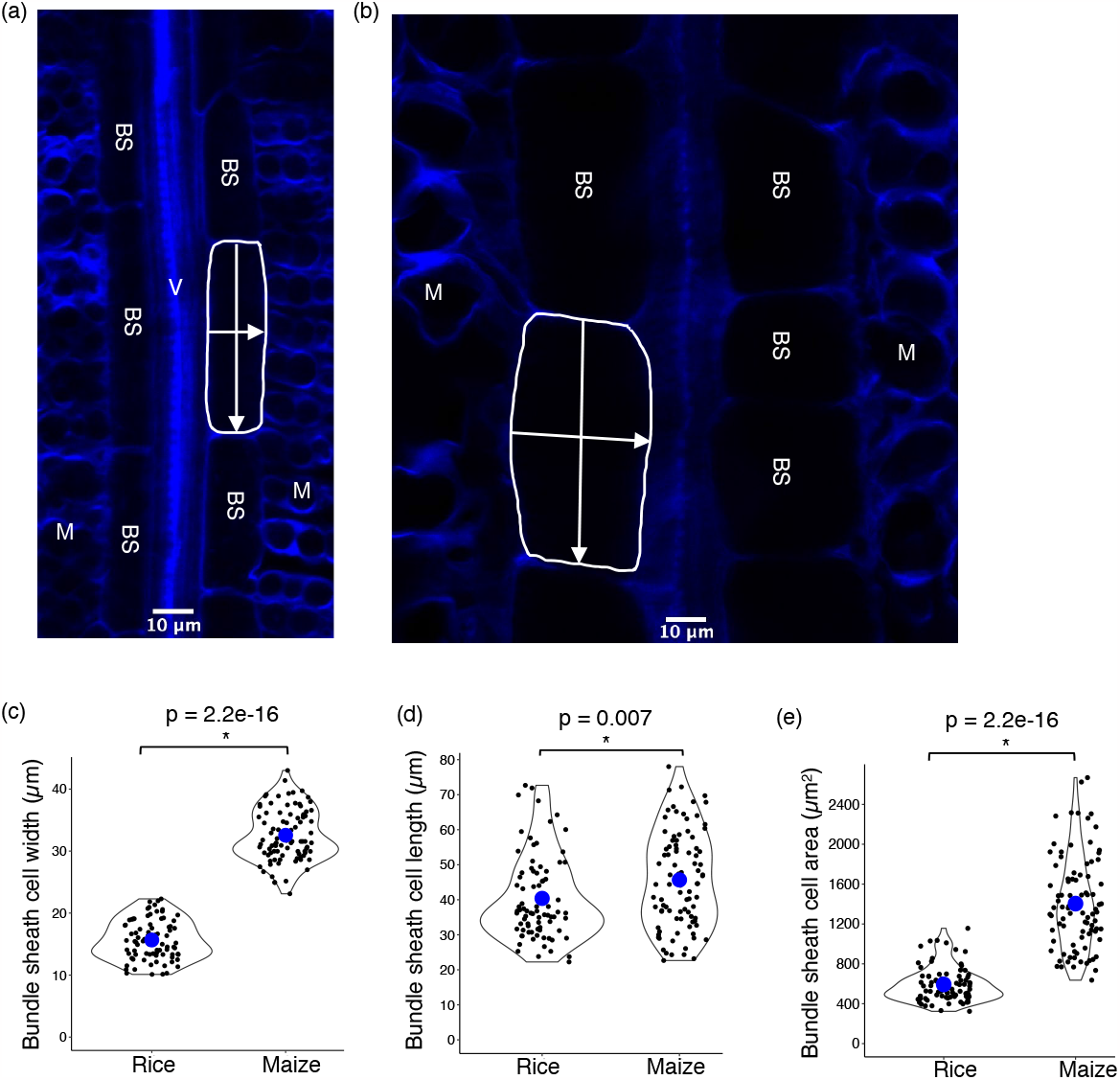
Quantification of bundle sheath cell dimensions in rice and maize. (a) Representative confocal laser scanning microscopy images of calcofluor white stained bundle sheath strand and mesophyll tissue of rice (a) or maize (b). Images were cropped to focus on a bundle sheath strand. Violin plots representing bundle sheath cell cell width (c), length (d) and area (e) in rice and maize. Blue dot represents mean values. Bundle sheath cell length and width measurements were taken at the mid-point of the proximal-distal and medio-lateral axes respectively (annotated with white arrows in a&b). Each observation represents one cell. Number of cells (n) = 82 and 90 from rice and maize, respectively. Abbreviations are as follows: V: Vein; BS: Bundle sheath cell; M: Mesophyll cell. Statistical test: t-test.

### Visualisation and quantification of chloroplast parameters in rice bundle sheath cells

We next wished to investigate whether bundle sheath chloroplast number and size could be determined after leaf ablation. Transgenic rice lines expressing the maize *GOLDEN2* (*ZmG2*) transcription factor under the control of the maize ubiquitin promoter are known to contain larger chloroplasts [22] and so were used as controls. We used calcofluor white to stain cell walls and chlorophyll autofluorescence to visualize bundle sheath cell chloroplasts. Z-stacks of 82 and 90 bundle sheath cells from wild-type and p*ZmUbi*::*ZmG2* rice respectively were acquired by confocal laser scanning microscopy. Maximum intensity projection images (Figure 4a) were used to quantify individual chloroplast areas and chloroplast number per cells. The average planar area of individual bundle sheath chloroplasts in wild-type was ∼ 17 μm^2^ (with a range from 7 to 37 μm^2^; Figure 4b). Consistent with published data [22] planar area of bundle sheath chloroplasts was significantly increased in the p*ZmUbi*::*ZmG2* line and ranged from 7 to 54 μm^2^ (Figure 4b). Moreover, as expected [22] there was no difference in chloroplast numbers in the bundle sheath between controls and p*ZmUbi*::*ZmG2* (Figure 4c). However, total chloroplast occupancy of bundle sheath cells in p*ZmUbi*::*ZmG2* was significantly increased due to the greater planar area of individual chloroplasts (Figure 4d). Further, total chloroplast number per bundle sheath cell (with a range from 8 to 25) obtained from leaf ablation (Figure 4c) was comparable with that previously reported from analysis of isolated single-cells (with a range from 6 to 21; [22]). Therefore, gentle and careful ablation can be used to obtain accurate estimates of chloroplast numbers in rice bundle sheath cells.

**Figure 4:**
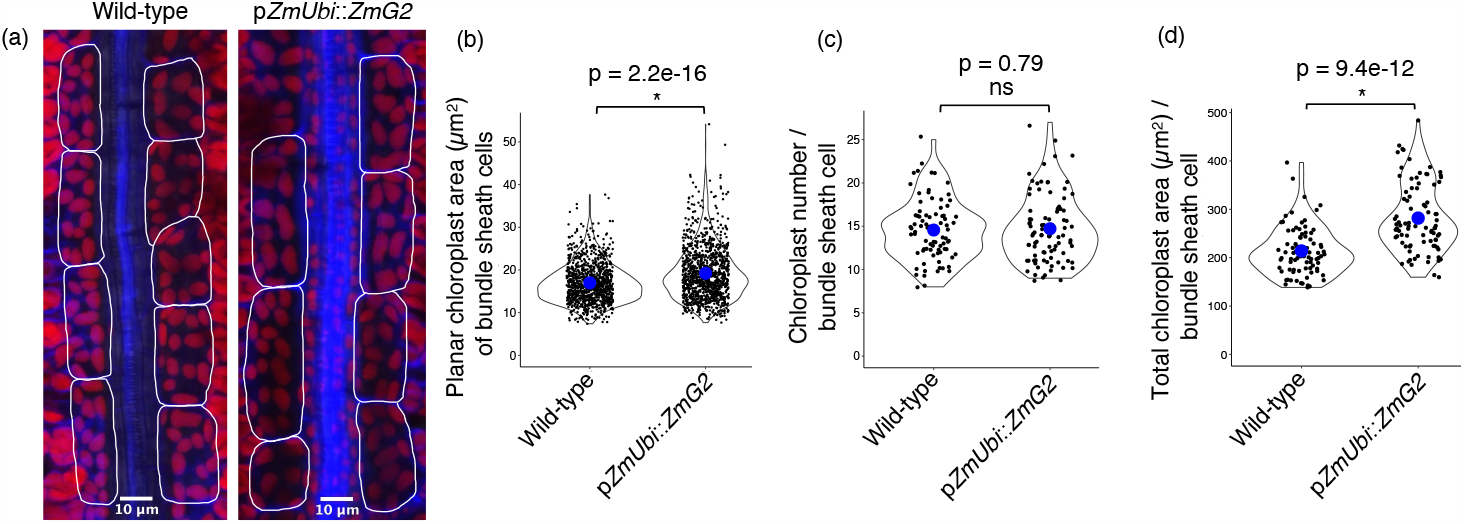
Visualization and quantification of chloroplast area and number in the bundle sheath of rice. (a) Representative maximum intensity projection image of a Z-stack from wild-type and the *GOLDEN2* overexpressing line (p*ZmUbi*::*ZmG2*). Images were cropped to focus on a bundle sheath strand. Bundle sheath cells are highlighted with white lines. (b) Planar area of individual chloroplasts from bundle sheath cells. Each observation represents one chloroplast. Number of chloroplasts (n) = 1032 and 1114 from wild-type and p*ZmUbi*::*ZmG2* rice lines, respectively. (c) Chloroplast number per bundle sheath cell. Each observation represents one cell. (d) Total chloroplast area per bundle sheath cell. Each observation represents the total chloroplast area of a cell. Number of cells (n) = 82 and 90 from wild-type and p*ZmUbi*::*ZmG2* rice lines, respectively. Blue dot in the violin plots represent mean values. Statistical test: t-test.

Acquisition of three-dimensional (3D) images is of course more time consuming than two-dimensional (2D) images. We therefore wanted to test if there was a difference between bundle sheath chloroplast numbers estimated by the two approaches and so obtained 2D and 3D images of the same 31 cells from wild-type (Figure 5a). These data showed that the bundle sheath chloroplast number was significantly higher (Figure 5b) when estimated from 3D imaging (with a range from 11 to 25) compared with 2D imaging (with a range from 10 to 19). However, planar area of individual chloroplasts in bundle sheath cells was not different between the two datasets (Figure 5c). We conclude that 3D imaging provides a more precise estimate of bundle sheath chloroplast numbers but either method can be used to quantify chloroplast size.

**Figure 5:**
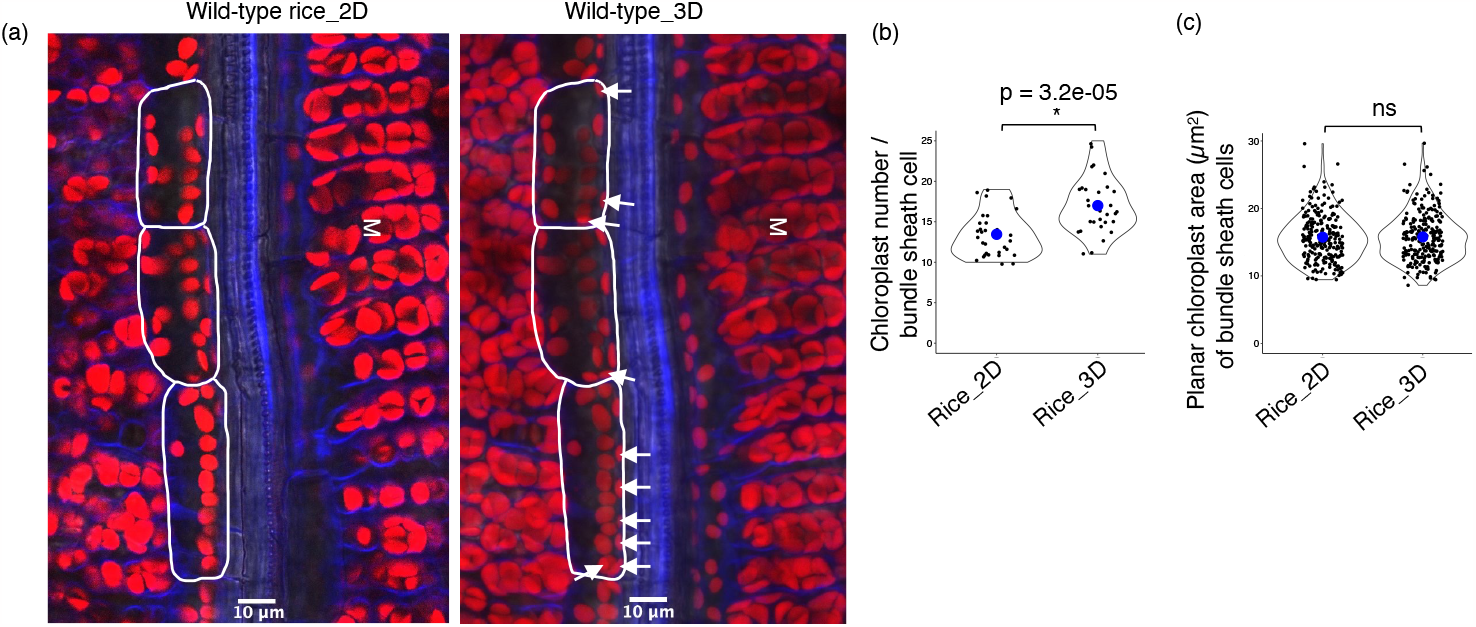
Comparison of chloroplast numbers obtained from two and three-dimensional (2D and 3D) imaging. (a) Representative two-dimensional (left) and three-dimensional (right) images of wild-type rice leaves. Bundle sheath cells are highlighted with white lines. Chloroplasts present in the 3D image but not detected in the 2D image are highlighted with white arrows. (b) Chloroplast number per bundle sheath cell - each observation represents one cell (n = 31). (c) Planar chloroplast area of bundle sheath cells - each observation represents one chloroplast. Number of chloroplasts (n) = 236 and 242 from 2D and 3D imaging, respectively. Blue dot in the violin plots represent mean values. Statistical test: t-test. M: Mesophyll tissue.

### The relationship between bundle sheath paradermal cell area and chloroplasts

We wanted to use the above data to understand the relationship between bundle sheath chloroplast occupancy and cell area in rice. Therefore, a simple linear regression model was performed between bundle sheath paradermal cell area and chloroplast size and number. This showed that the average planar and maximum chloroplast area per cell did not vary with bundle sheath cell area (Figure 6a&b). But, chloroplast number and thus total chloroplast area per cell increased with cell area (Figure 6c and d). The percentage of cell area occupied by chloroplast was negatively correlated with bundle sheath cell area (Figure 6e).

**Figure 6:**
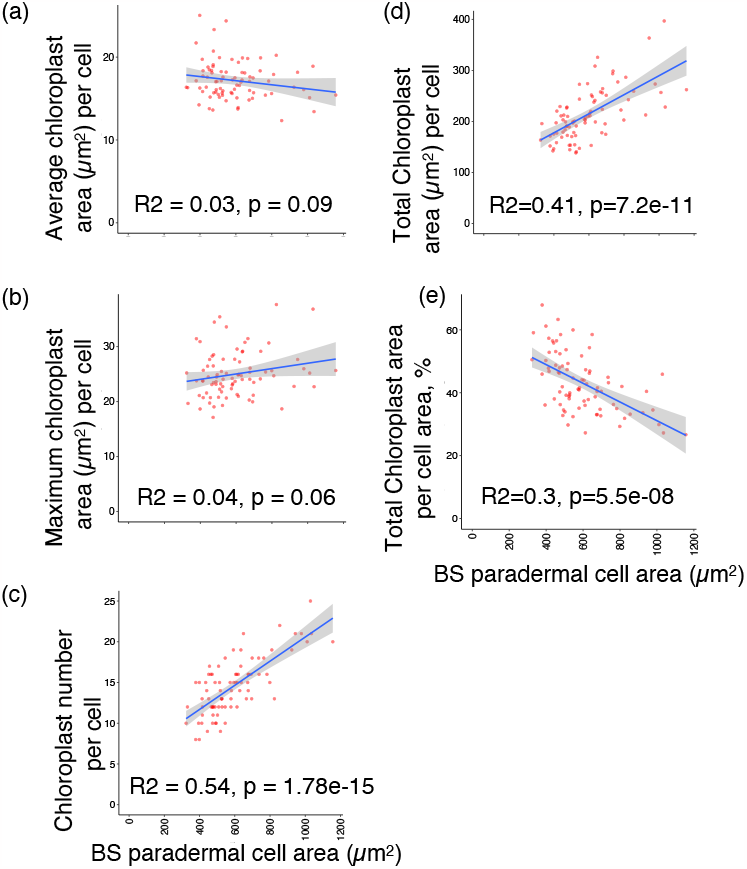
Linear regression analysis between bundle sheath chloroplast parameters and paradermal cell area. Linear regression plots representing bundle sheath (BS) paradermal cell area versus average chloroplast area (a), maximum chloroplast area (b), chloroplast numbers (c), total chloroplast occupancy (d), and relative chloroplast area in the cell (%). Graphs show the line of best fit and standard error (gray filled region) of the linear model fitted to the data (red circles).

## Discussion

It is widely recognised that improving photosynthesis in crops is one mechanism to improve yield [27]. One approach that has been proposed [17,28] is to engineer the C_4_ pathway into C_3_ crops such as rice and it is estimated that this could improve yields by up to 50 %. However, this goal is challenging and would require a significant increase in the chloroplast compartment of bundle sheath cells from C_3_ crops such as rice. It has been challenging to phenotype bundle sheath tissue in C_3_ species as these cells are deeper in the leaf because of the many layers of mesophyll cells [25]. Approaches including bright-field light microscopy [18], transmission electron microscopy [23], serial block-face scanning electron microscopy [24] and single-cell isolation methods [22] are slow and so this hinders rapid analysis of transgenic lines harbouring candidate genes that are hypothesized to control chloroplast proliferation in the bundle sheath. To this end, we sought to establish a rapid and robust method to visualize bundle sheath cell files in C_3_ rice.

Including sample preparation time, the ablation method reported here requires about 30 minutes to phenotype one leaf sample and can capture images from 30-40 bundle sheath cells in one focal plane. To obtain three-dimensional imaging via acquisition of z-stacks approximately one hour is needed. This compares favourably with other approaches such as the published single-cell isolation method [22] which involves five hours of sample preparation followed by three-four hours to image a similar number of cells. Thus, we estimate that the leaf ablation method is at least ten times faster than single-cell isolation.

Other methods that involve resin-embedding, thin-sectioning, and then image capture via light or electron microscopy take a few weeks. The leaf ablation method also excludes hazardous chemicals and enzymes for sample preparation, and it is noteworthy that it also allows specific vein types to be identified prior to imaging, which can be challenging with the single-cell isolation method as the leaf tissue is subject to enzymatic digestion. We therefore consider this simple ablation approach to be robust and useful for high-throughput *in vivo* phenotyping of bundle sheath cells in C_3_ species.

To provide evidence that imaging after ablation captures parameters derived from intact bundle sheath cells, the width of rice bundle sheath cells was measured from transverse sections (Additional file 2a) and compared with paradermal cell width obtained from confocal microscopy imaging after leaf ablation (Additional file 2b). As bundle sheath cells are cylindrical, the width should equal the depth. In fact, mean bundle sheath cell width was lower (∼10 μm) when estimated from transverse sections compared with paradermal sections (∼15 μm; Additional file 2b) implying that the estimates of cell width after ablation are not associated with incomplete imaging of this cell type. It is also possible that paradermal sections preferentially captured information on bundle sheath cells lateral to each vein (Figure 3a). It has been reported that during the C_3_ to C_4_ trajectory bundle sheath cells elongate less along the axis of the vein but become wider [14,15]. Consistent with this, we observed a two-fold increase in bundle sheath cell width in maize compared with rice (Figure 3c). However, the length of bundle sheath cells in the two species were similar and so these data suggest that a reduction in bundle sheath cell length may not be required for the evolution of C_4_ photosynthesis.

Under the conditions we used the average planar area of bundle sheath chloroplasts in wild-type and p*ZmUbi*::*ZmG2* were higher than in previous analysis [22]. These differences might result from different experimental conditions and/or from the use of confocal laser microscopy to study chloroplasts in our study. For example, Wang et al., 2017 used the single-cell isolation method followed by bright-field light microscopy. There is a possibility that chloroplast area is over-estimated from chlorophyll autofluorescence due to the introduction of background pixels. To investigate this, we measured the planar area of 574 bundle sheath chloroplasts from the two-dimensional images of wild-type rice leaf samples obtained from serial block-face scanning electron microscopy (Additional file 3a). Based on this approach, the average area of individual chloroplasts was 14 μm^2^ (Additional file 3b), which is higher than previously reported (11 μm^2^; [22]) but lower than what we estimated from confocal imaging after ablation (16-17 μm^2^; Figure 4b & 5c). Thus, it implies that the larger chloroplast area might result from both our experimental conditions and confocal imaging. Irrespective of these differences, leaf ablation in association with confocal imaging allows differences between genotypes to be detected (Figure 4b&d).

## Conclusions

In conclusion, we report a simple and scalable leaf ablation method to access bundle sheath cell files in C_3_ species such as rice. We show that this method is appropriate to measure bundle sheath cell dimensions, chloroplast areas and chloroplast numbers per cell. We also show bundle sheath cells are intact after the leaf ablation. As the approach is at least ten times faster than the next most efficient approach, ablation should significantly accelerate analysis of transgenic lines harbouring candidate genes aimed at modifying the rice bundle sheath.

## Materials and methods

### Plant material and growth conditions

Seeds of wild-type (*Oryza sativa spp japonica* cv. Kitaake) and maize *GOLDEN-2* (*ZmG2*) overexpressing rice ([22]; *ZmUBI*_pro_::*ZmG2* line E131) were imbibed in sterile Milli-Q water and incubated at 30°C in the dark for two days. Seeds were transferred onto Petri plates with moistened Whatman filter paper and germinated in the growth cabinet at 28°C with 16/8 hrs. of light/dark cycle. After two days, germinated seedlings were potted into 9 by 9 cm pots (two plants/pot) filled with Profile Field and Fairway soil amendment (www.rigbytaylor.com). Plants were grown in a walk-in plant growth chamber under a 12-hour photoperiod at a photon flux density of 400 μmol m^-2^ sec^-1^ at 28°c (day) and 20°C night. Once a week, plants were fed with the Peters Excel Cal-Mag Grower fertiliser solution (LBS Horticulture, Clone, UK) with additionally supplied iron (Fe7 EDDHA regular, Gardening Direct, UK). The working fertiliser solution contains 0.33 g/L of Peters Excel Cal-Mag Grower and 0.065 g/L chelated iron. Maize (B73) was grown in a growth cabinet operating at 28°C (day)/ 20°C (night) at a photon flux density of 550 μmol m^-2^ sec^-1^ under a 14-hour photoperiod.

### Sample preparation

The middle region of the fully expanded fourth leaf from wild-type Kitaake, *ZmUBI*_pro_::*ZmG2* overexpressing rice lines and maize was fixed with 1 % (w/v) glutaraldehyde in 1X PBS buffer. Once fixative was infiltrated, samples were left in that solution for about two hours and then washed twice with 1X PBS buffer, with each wash lasting ∼ 30 minutes. Leaf samples can be stored in 1X PBS buffer at 4°C for several weeks without losing chlorophyll autofluorescence. Before microscopy, the adaxial side of the fixed leaf material was ablated gently with a fine razor blade (Personna, Verona, VA 24482) to remove mesophyll layers. Bundle sheath cells can be directly visualized with light microscopy. For confocal microscopy, the ablated leaf fragment was stained with the cell wall stain calcofluor white (0.1 %; Sigma) for 5 mins and then rinsed twice with H_2_O.

### Light and confocal laser microscopy

Light microscopy images (Olympus BX51 microscope) were captured using an MP3.3-RTV-R-CLR-10-C MicroPublisher camera and QCapture Pro 7 software (Teledyne Photometrics, Birmingham, UK). A Leica SP8X confocal microscope upright system (Leica Microsystems) was used for fluorescence imaging. It has two continuous wave laser lines, 405 nm and 442 nm, a 460-670 nm super continuum white light laser (WLL) and four hybrid detectors and one photomultiplier tube. Imaging was conducted using a 25X water immersion objective and Leica Application Suite X (LAS X; version: 3.5.7.23225) software. Calcofluor white was excited at 405 nm and emitted fluorescence captured from 452–472 nm. Chlorophyll autofluorescence was excited at 488 nm and emission captured 672–692 nm. Three replicates from both wild-type Kitaake and *ZmUBI*_pro_::*ZmG2* overexpression line E131 were analysed. Z-stacks of ∼ 30 lateral bundle sheath cells surrounding three different intermediate veins (3°) and eight to ten cells per vein were obtained from each replicate. From three replicates, 82 and 90 bundle sheath cells from wild-type and E131 line were imaged respectively. Images of 90 maize bundle sheath cells were captured using light microscopy to measure bundle sheath cell dimensions.

### Serial block-face scanning electron microscopy

Wild-type rice leaf (middle region of fourth leaves) samples were fixed in fixative (2 % w/v glutaraldehyde / 2 % w/v formaldehyde in 0.05 M sodium cacodylate buffer pH 7.4 containing 2 mM calcium chloride) overnight at 4°C. After washing five times with 0.05 M sodium cacodylate buffer pH 7.4, samples were osmicated (1 % osmium tetroxide, 1.5 % potassium ferricyanide, 0.05 M sodium cacodylate buffer pH 7.4) for three days at 4°C.

After washing five times in DIW (deionised water) samples were treated with 0.1 % (w/v) thiocarbohydrazide/DIW for 20 minutes at room temperature in the dark. After washing five times in DIW, samples were osmicated a second time for one hour at RT (2 % osmium tetroxide/DIW). After washing five times in DIW, samples were block stained with uranyl acetate (2 % uranyl acetate in 0.05 M maleate buffer pH 5.5) for three days at 4°C. Samples were washed five times in DIW and then dehydrated in a graded series of ethanol (50 %/70 %/95 %/100 %/100 % dry), 100 % dry acetone and 100 % dry acetonitrile, three times in each for at least five minutes. Samples were infiltrated with a 50/50 mixture of 100 % dry acetonitrile/Quetol resin mix (without BDMA) overnight, followed by three days in 100 % Quetol (without BDMA). Then, the sample was infiltrated for five days in 100 % Quetol resin with BDMA, exchanging the resin each day. The Quetol resin mixture is: 12 g Quetol 651, 15.7 g NSA (nonenyl succinic anhydride), 5.7 g MNA (methyl nadic anhydride) and 0.5 g BDMA (benzyldimethylamine; all from TAAB). Samples were placed in embedding moulds and cured at 60°C for three days.

Sections were cut at a thickness of about 70 nm using a Leica Ultracut E, placed on a Melinex plastic coverslip, and allowed to air dry. Coverslips were mounted on aluminium scanning electron microscopy stubs using conductive carbon tabs and the edges of the slides were painted with conductive silver paint. Then, samples were sputter coated with 30 nm carbon using a Quorum Q150 TE carbon coater. Samples were imaged in a Verios 460 scanning electron microscope (FEI/Thermofisher) at 4 keV accelerating voltage and 0.2 nA probe current in backscatter mode using the concentric backscatter detector (CBS) in field-free mode for low magnification imaging and in immersion mode at a working distance of 3.5-4 mm; 1536×1024 pixel resolution, 3 us dwell time, 4 line integrations for higher magnification imaging. Stitched maps were acquired using FEI MAPS automated acquisition software using the default stitching profile and 5 % image overlap. Transverse bundle sheath cell width was measured from bundle sheath cells of three minor veins per replicate, and three biological replicates were used. In total, dimensions of 92 bundle sheath cells were measured. The planar chloroplast areas were measured from paradermal sections of bundle sheath cells surrounding two minor veins per replicate. Total areas of 574 chloroplasts were measured across 130 cells.

### Data analyses

Bundle sheath cell dimensions (length, width, and area), chloroplast area and numbers per cell were measured using ImageJ version 2.1.0/1.53c [29]. RStudio (version:1.4.1106) was used to plot the data using the ggplot2 software package [30] and statistical analysis was performed using the ggpubr software package [31]. First, equality of variance between the two groups was tested using Barlett’s test [32]. Where the assumption of equal variance was met, a two-tailed pairwise t-test (Student’s t-test) was performed. Otherwise, Welch’s two-sample t-test was performed. A general linear regression model was performed using the ggfortify package [33] and assumptions of a linear regression model were tested using the autoplot function of the ggfortify package. Finally, the general linear regression line was fitted using the lm function and, ANOVA test was performed to test whether the slope is significantly different from zero.

## Additional information

**Additional file 1:** Movie showing rice leaf ablation.

Rice leaf image with different vein orders before and after leaf ablation (top) and video showing the leaf ablation process (bottom). A drop of water was added onto a glass plate to prevent the dehydration while ablating the leaf. 1°: primary/mid vein; 2°: secondary/large lateral veins; 3°: tertiary/intermediate veins. Movie courtesy: Dr Satish Kumar Eeda.

**Additional file 2:** Comparison of bundle cell width in paradermal versus transverse sections.

(a) Transverse section of a rice leaf obtained from serial block-face scanning electron microscopy, representing the bundle sheath cells of a tertiary vein (3°). Bundle sheath cell width was measured at the mid-point of the medio-lateral axes as annotated with a red arrow. (b) Comparison of bundle sheath cell width measurements from paradermal and transverse sections, obtained from confocal imaging (rice data from Figure 3c) and serial block-face scanning electron microscopy, respectively. BS: Bundle sheath cell; M: Mesophyll cell. Blue dot in the violin plots represent mean values. Statistical test: t-test.

**Additional file 3:** Comparison of individual chloroplast areas obtained from confocal laser scanning microscopy versus serial block-face scanning electron microscopy (SBF-SEM).

(a) Paradermal section of a rice leaf obtained from serial block-face scanning electron microscopy, representing the lateral bundle sheath cells of a tertiary vein (3°). Bundle sheath chloroplasts were pointed with red arrows. (b) Comparison of individual chloroplast areas from confocal (wild-type rice data from Figure 4b) and two-dimensional serial block-face scanning electron microscopy imaging. BS: Bundle sheath cell; M: Mesophyll cell. Blue dot in the violin plots represent mean values. Statistical test: t-test.

## Acknowledgements

This research was funded by a C_4_ Rice Project grant (#INV-002970) from The Bill & Melinda Gates Foundation to the University of Oxford. For the purposes of open access, the authors have applied a Creative Commons Attribution (CC BY) license to any Author Accepted Manuscript version arising from this submission. We thank Lei Hua for useful suggestions for confocal laser microscopy work, Lee Cackett for providing maize leaf material and Tina B. Schreier for assistance with serial block-face scanning electron microscopy imaging. We also thank Jane Langdale for providing rice seeds of *ZmUBI*_pro_::*ZmG2* overexpressing line. We thank Karin H Müller and Georgina E Lindop from the Cambridge Advanced Imaging Centre (CAIC) for the electron microscopy sample preparation as well as image acquisition.

## Author contributions

KB designed and performed the experiments. KB drafted the manuscript. KB and JMH reviewed and edited the manuscript.

## Declarations

### Ethics approval and consent to participate

Not applicable

### Consent for publication

Not applicable

### Availability of data and materials

All data supporting the findings of this study are available within the paper and within its supporting information data published online.

