## Additional file 2 for "A rapid and robust leaf ablation method to visualize bundle sheath cell chloroplasts in C_3_ species"

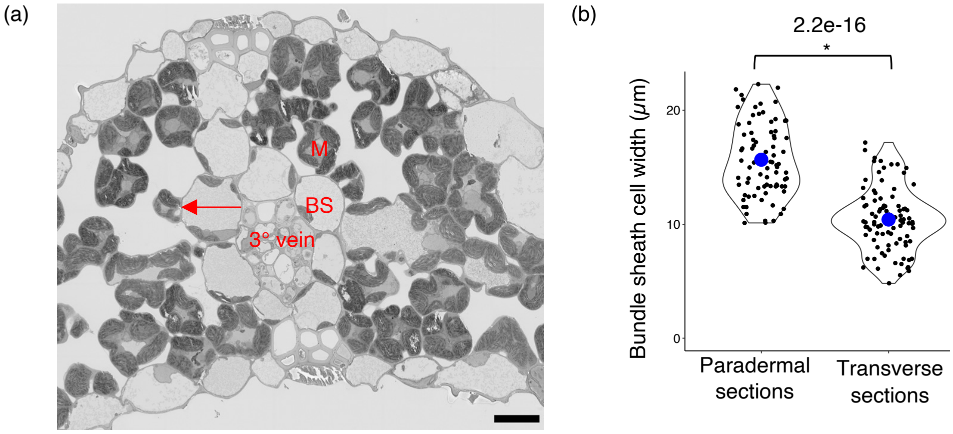


**Additional file 2:** Comparison of bundle cell width in paradermal versus transverse sections

(a) Transverse section of a rice leaf obtained from serial block-face scanning electron microscopy, representing the bundle sheath cells of a tertiary vein (3°). Bundle sheath cell width was measured at the mid-point of the medio-lateral axes as annotated with a red arrow. (b) Comparison of bundle sheath cell width measurements from paradermal and transverse sections, obtained from confocal imaging (rice data from Figure 3c) and serial block-face scanning electron microscopy, respectively. BS: Bundle sheath cell; M: Mesophyll cell. Blue dot in the violin plots represent mean values. Statistical test: t-test.
