## Additional file 3 for "A rapid and robust leaf ablation method to visualize bundle sheath cell chloroplasts in C_3_ species"

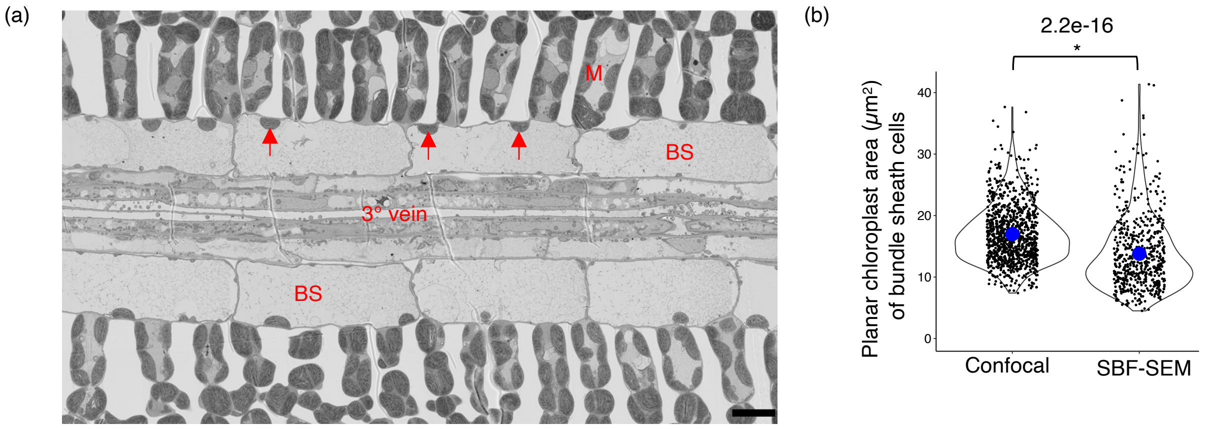


**Additional file 3:** Comparison of individual chloroplast areas obtained from confocal laser scanning microscopy versus serial block-face scanning electron microscopy (SBF-SEM)

(a) Paradermal section of a rice leaf obtained from serial block-face scanning electron microscopy, representing the lateral bundle sheath cells of a tertiary vein (3°). Bundle sheath chloroplasts were pointed with red arrows. (b) Comparison of individual chloroplast areas from confocal (wild-type rice data from Figure 4b) and two-dimensional serial block-face scanning electron microscopy imaging. BS: Bundle sheath cell; M: Mesophyll cell. Blue dot in the violin plots represent mean values. Statistical test: t-test.
