## Additional file 1 for "A rapid and robust leaf ablation method to visualize bundle sheath cell chloroplasts in C_3_ species"

### Slide 1
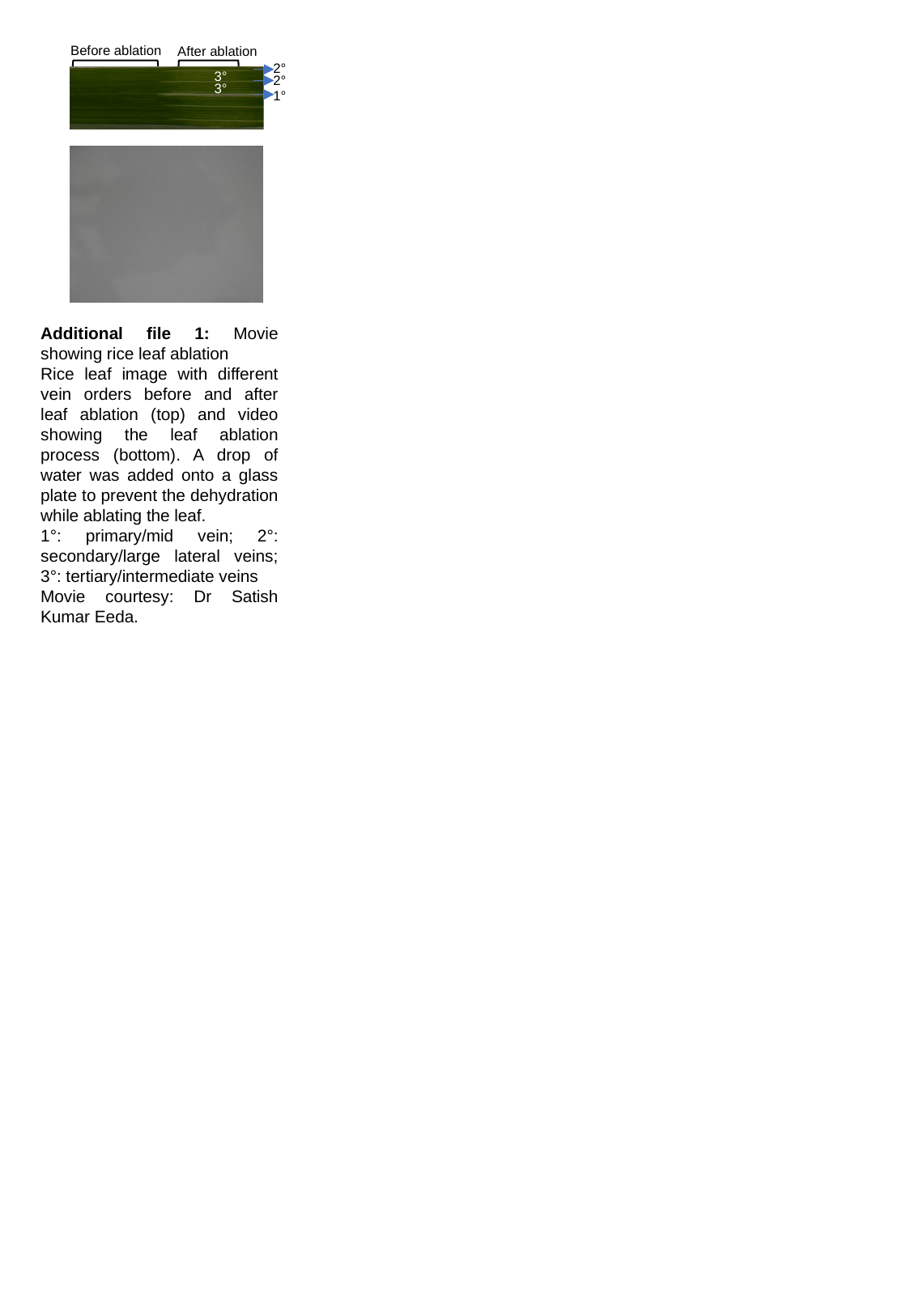

Before ablation
After ablation
2°
3°
2°
3°
1°
Additional file 1: Movie showing rice leaf ablation
Rice leaf image with different vein orders before and after leaf ablation (top) and video showing the leaf ablation process (bottom). A drop of water was added onto a glass plate to prevent the dehydration while ablating the leaf.
1°: primary/mid vein; 2°: secondary/large lateral veins; 3°: tertiary/intermediate veins
Movie courtesy: Dr Satish Kumar Eeda.
